# A fine-scale genetic map of the Japanese population

**DOI:** 10.1101/2023.09.19.558557

**Authors:** Jun Takayama, Satoshi Makino, Takamitsu Funayama, Masao Ueki, Akira Narita, Keiko Murakami, Masatsugu Orui, Mami Ishikuro, Taku Obara, the Tohoku Medical Megabank Project Study Group, Shinichi Kuriyama, Masayuki Yamamoto, Gen Tamiya

## Abstract

Genetic maps are fundamental resources for linkage and association studies. A fine-scale genetic map can be constructed by inferring historical recombination events from the genome-wide structure of linkage disequilibrium—a non-random association of alleles among loci—by using population-scale sequencing data. We constructed a fine-scale genetic map and identified recombination hotspots from 10,092,573 bi-allelic high-quality autosomal markers segregating among 150 unrelated Japanese individuals. These individuals’ genotypes were determined by high-coverage (30×) whole-genome sequencing, and the genotype quality was carefully controlled by using their parents’ and offspring’s genotypes. The pedigree information was also utilized for haplotype phasing. The resulting genome-wide recombination rate profiles were concordant with those of the HapMap worldwide population on a broad scale, and the resolution was much improved. We identified 9487 recombination hotspots and confirmed the enrichment of previously known motifs in the hotspots. Moreover, we demonstrated that the Japanese genetic map improved the haplotype phasing and genotype imputation accuracy for the Japanese population. The construction of a population-specific genetic map will help make genetics research more accurate.

## INTRODUCTION

Genetic maps play a pivotal role in human genetic studies [1–3], including association studies and gene mapping through linkage analysis. They provide relative distances between genetic markers on chromosomes in terms of meiotic recombination rates. Meiotic recombination events do not occur evenly across chromosomes; rather, their occurrences tend to be concentrated in relatively restricted, narrow regions (so-called recombination hotspots [4]), creating block-like structures called haplotype blocks [5–9]. Genetic maps can be constructed by estimating recombination events in families, sperm typing, radiation hybrid mapping, or inferring linkage disequilibrium (LD) through population genotyping [10–12]. LD refers to the non-random association between alleles at different loci. The LD structure across the genome is shaped by population genetics forces such as recombination, genetic drift, mutation, selection, and gene conversion [13]. Here, we estimated the population-scale recombination rate from the LD structure by assuming an equilibrium between recombination and genetic drift. Thus, by inspecting the LD structure in a population, we inferred historical recombination events and constructed a genetic map.

The recent advent of next-generation sequencing technologies has enabled population-scale genotyping, which can be used to genotype millions of single nucleotide variant (SNV) markers for thousands of individuals and thus construct fine-scale genetic maps [11, 12]. However, because the inferred LD structure is substantially distorted by mutations and erroneous genotypes mimicking recombination events, quality control (QC) of the genetic markers is critical. Moreover, because of differences in population demography, LD structure differs substantially among populations [7, 13, 14]: for example, the African population has shorter LD blocks and denser recombination hotspots, whereas European and Asian populations have longer LD blocks and sparser hotspots.

Large-scale human population genetic studies have so far been used to construct LD-based fine-scale genetic maps. The HapMap project has constructed a genetic map from three worldwide populations: Africans, Asians (Japanese + Chinese), and Europeans [15]. As well as the standard marker QC procedures applicable to unrelated individuals, including Hardy-Weinberg equilibrium tests, the HapMap genetic map markers were examined for Mendelian errors in a QC step that utilized pedigree information from 30 European and 30 African trio families. However, no Asian families were collected in the project. Hence the 90 Asian samples were genotyped at the single nucleotide polymorphism (SNP) markers that passed the Mendelian error QC of the African and European family samples. The 1000 Genomes Project (1KGP) has also constructed an LD-based genetic map for worldwide populations [16]. However, the 1KGP genetic map was constructed solely by using unrelated individuals and did not utilize pedigree information for QC. Moreover, the 1KGP samples were sequenced at low coverage (7.4×). Consequently, there are no fine-scale Japanese genetic maps for which the marker quality has been ascertained by using pedigree information. Therefore, the creation of a fine-scale genetic map for the Japanese population by using pedigree-based QC is critical for inferring historical recombination events among entire human populations and for performing genetic analysis of Japanese populations.

In addition to accurate genotyping, inference of LD structure requires accurate haplotype phasing information. Although haplotype phasing is possible without pedigree information, the pedigree information provides more accurate phasing information. Therefore, it is preferable to use pedigree information for haplotype phasing, if available.

Here, we constructed a fine-scale, high-quality, LD-based genetic map for the Japanese population from 10 million SNPs of 150 deeply sequenced unrelated individuals; genotype QC and haplotype phasing were performed by using genotype information from their parents and offspring. The broadscale recombination rate variation of the Japanese population resembled that of the HapMap population, and the resolution of the fine-scale rate variation was much improved. We identified recombination hotspots and confirmed the enrichment of known sequence characteristics in the hotspots. We also demonstrate the usefulness of the population-specific genetic map.

## MATERIALS AND METHODS

### Sample selection

We selected 1282 Japanese individuals with whole-genome sequencing (WGS) data from the Tohoku Medical Megabank Project (TMM) BirThree cohort study, a prospective genomic cohort of individuals including family information in the Tohoku area in Japan [17, 18]. We excluded individuals for whom the WGS-based family relationships and recorded relationships were inconsistent. We then excluded an eight-member family that had the highest number of Mendelian errors. We also excluded 132 individuals belonging to families with low read-depth data, as analyzed by using saliva-derived DNA. This filtering retained 1120 individuals, which were then subject to marker QC. Among the 1120 individuals, we restricted the downstream analyses to 773 individuals analyzed by Illumina NovaSeq 6000 sequencer. Among these 773 individuals, we further excluded eight who appeared to belong to the Chinese population according to our principal component analysis. We also excluded a seven-member family, in which a member was in cousin relationships with other individuals in the dataset. The remaining 758 individuals consisted of 23 eight-member three-generation families (184 individuals) and 82 seven-member three-generation families (574 individuals). We selected all 23 eight-member families and 52 seven-member families with the least Mendelian errors (**Supplementary Fig. 1**). From these 75 families, we subjected the 150 individuals of generation II (i.e., fathers and mothers) to the subsequent analysis.

### Marker QC

Marker QC was performed in accordance with the method of Auton et al. [12] where possible. We excluded the following regions: (1) regions with three or more SNPs within 10 bp; (2) sites fixed to non-reference alleles for all individuals; (3) sites with quality score divided by the sum of read depth of individuals with non-reference genotypes ≥ 27.0 or ≤ 1.0; and (4) those with an estimated Phred-scaled probability of ≥ 40 that read bases within 10 bp did not have more than two haplotypes. After applying these filters, we extracted bi-allelic sites. We then defined a cohort of the 1120 individuals mentioned in the “Sample selection” section above and 3315 other unrelated individuals from the cohort used to build the 3.5KJPNv2 allele frequency panel of Japanese individuals [18]. This resulted in a total of 4435 individuals. We then excluded (1) insertion-deletion (indel) sites; (2) SNPs on indel sites; (3) SNP sites with a missing rate > 5% (also applied in the work of Auton et al. [12]); (4) in-house low-quality markers; (5) singletons; and (6) sites with a *P*-value < 0.01 in the Hardy-Weinberg equilibrium test. After this filtering, we obtained 23,160,390 bi-allelic SNPs across the 1120 individuals for the subsequent analyses.

We phased the quality-controlled 23,160,390 SNP sites by using SHAPEIT2+duoHMM (ver. 2.900) software [19], which can improve phasing by using pedigree information. For phasing, we set the window size to 5 Mb, and the number of hidden Markov model states to 300. In addition, we used the 1KGP genetic map as the initial value. The resulting dataset consisted of 10,092,573 variants and 300 haplotypes.

### Construction of an LD-based genetic map

We estimated the population-scale historical recombination rates by using LDhat/interval (ver. 2.2) software [10, 11]. We estimated the rates in chunks of 4000 sites, each with a 200-site overlap. The block penalty to avoid overfitting was set to 5. Rates were estimated by sampling once every 15,000 iterations out of 30,300,000 iterations. We used LDhat/complete software to generate a likelihood lookup table file for 300 haploids in 900 chunks, assuming that the population mutation rate *θ* = 0.001 [20]. Summary statistics were obtained by using LDhat/stat software. We set the per-kilobase mean population recombination rate, *ρ*, to 0 for regions with *ρ >* 100. We also set *ρ* to 0 for regions containing >50 kb gaps and their 50 upstream and 50 downstream sites. We converted our *ρ* estimates into per-generation recombination rates, *r*, given the effective population size *N*_e_ (*ρ* = 4*N*_e_*r*), by linear regression on the deCODE genetic map [3] with the lm function in R (ver. 3.5.1) software.

### Identification of recombination hotspots

We identified recombination hotspots by using LDhot software (ver. 0.4) [21] from 1000 simulations, setting the background window and the hotspot window to ± 100 kb and ± 1 kb of the hotspot center, respectively. Post-processing was performed by using LDhot summary software. We excluded candidate hotspots with peak rate estimates < 5 or those with width ≥ 5 kb to reduce false positives according to ref. [12].

### Hotspot enrichment analysis

Control sets with matched lengths selected from matched chromosomes were prepared by using BEDTools (ver. 2.29.1) software [22]. Interspersed and tandem repeats were identified by RepeatMasker (ver. 4.0.7) software [23]. GC% was obtained by using the seqtk (ver. 1.3) software comp command. Motif enrichment was assessed by using AME (ver. 5.0.2) software [24] of the MEME suite by comparison with the control regions described above. We obtained MEME-format motif files by using the iupac2meme command of the MEME suite, providing CCTCCCT and CCNCCNTNNCCNC as input for the 7-bp and 13-bp motifs, respectively. Local motif enrichment within the hotspots was assessed by using CENTRIMO (ver. 5.0.2) software [25] of the MEME suite. Because CENTRIMO input sequences must be of the same size, we obtained the central 3000 bp by using fasta-center software of the MEME suite. CENTRIMO was run with the control regions as the negative control.

### Haplotype phasing to evaluate the utility of our genetic map

To assess the accuracy of phasing with our genetic map, we used the WGS data of 104 individuals from the generation II of the TMM BirThree cohort, all of which were generated by Illumina HiSeq2500 sequencer. Markers on chromosome 10 were used for the assessment. Marker QC was performed as follows; sites with Mendelian error were set as missing, sites with MAF ≥ 0.01, call rate ≥ 0.99, and *P*-value for HWE ≥ 0.00001 were included. The haplotype phasing was performed by using SHAPEIT2 v2.904 [25], with varying *S* and *W* parameters; *S* was chosen from 100, 200, and 400, and *W* was chosen from 0.1, 0.5, 2, and 5. The switch error rate was calculated by using SHAPEIT5/switch_static v5.1.0 with their parents’ information [26].

### Genotype imputation to evaluate the utility of our genetic map

The genotype imputations with pre-phasing were performed by using SHAPEIT v2.900 [26] and IMPUTE2 v2.3.2 [28], with either our genetic map or the 1KGP genetic map, setting the effective population size to 20,000. We then compared the INFO scores (measures of imputation accuracy).

### HapMap public data

The HapMap genetic map [15] on the GRCh37 coordinate was downloaded from ftp://ftp.ncbi.nlm.nih.gov/hapmap/recombination/2011-01_phaseII_B37/genetic_map_HapMapII_GRCh37.tar.gz. Hotspot data on the NCBI35 coordinate system were downloaded from ftp://ftp.ncbi.nlm.nih.gov/hapmap/recombination/2006-10_rel21_phaseI+II/hotspots/hotspots.txt.gz. The hotspot regions were lifted over to the GRCh37 coordinate system by using liftOver software [29] with the hg17ToHg19.over.chain file, downloaded from http://hgdownload.soe.ucsc.edu/downloads.html.

## RESULTS

### Construction of an LD-based fine-scale genetic map for the Japanese population

To construct an LD-based fine-scale genetic map for the Japanese population, we used a dataset comprising 300 haploid genomes of 150 genetically unrelated Japanese individuals. The 150 individuals consisted of 75 males and 75 females in generation II (e.g., husbands and wives) from 75 three-generation families, which were recruited as part of a prospective genomic cohort [17] (**Supplementary Fig 1**). All family members from generations I to III were genotyped by WGS with a single sequencing platform, Illumina NovaSeq 6000, and jointly analyzed by using a single bioinformatics pipeline [18]. These uniform experimental and analytical conditions were expected to be free from platform bias. The average read depth per individual was 30.8 ± 3.0× (mean ± SD; n = 150 individuals), an appropriate depth for genome-wide genotyping without imputation [30].

Marker quality is critical for inferring LD structure, so we followed the stringent QC described by Auton et al. [12], where possible (see **Materials and Methods**). After applying this QC, we selected autosomal bi-allelic SNP sites that met the following criteria: the SNP (1) was not a singleton; (2) segregated among a cohort of 1120 individuals of the three-generation families and other cohort members consisting of 3315 unrelated Japanese individuals [18]; (3) was not identified as one of our in-house-defined low-quality markers; and (4) was not on short indel sites. These QC procedures produced 23,168,174 sites. After the QC application, we excluded 7784 sites with a ≥ 1% Mendelian error rate and set individual genotypes to missing for 365,393 sites with a <1% Mendelian error rate. Applying these filters resulted in 23,160,390 bi-allelic SNP sites among the 1120 individuals.

Next, we phased each allele on these approximately 23 million SNP sites on the basis of the direct relatives’ genotype information. This phasing procedure and selection of the 150 individuals mentioned above resulted in 10,092,573 phased bi-allelic autosomal SNP sites for the 150 individuals. Thus, we obtained a dataset of approximately 10 million bi-allelic SNP sites and 300 haploids.

We then assessed the characteristics and qualities of the 10 million SNP sites (**Supplementary Table 1**). The genome-wide marker SNP density was 3.76 sites/kb, which was adequate for detecting a typical hotspot size of 2 kb [31]. The average variant number per individual was 1,781,147 ± 8499 (mean ± SD; n = 150 individuals), and that per haploid genome was 1,093,017 ± 6550 (mean ± SD; n = 300 haploids). The genome-wide transition per transversion ratio of the 10 million SNP sites was 2.16, which is a generally observed value [32].

We then used LDhat/interval software [10, 11] to estimate *ρ* across the autosomes from the population genetic data described above. We scaled *ρ* to *r* by linear regression on the deCODE genetic map [3] in accordance with the method of Auton et al. [12]. *N*_e_ was 11,847, which was more similar to that of the European population (10,040) than that of the African population (19,064), as estimated by the same procedure [12]. In this way, we constructed a fine-scale genetic map for the Japanese population.

### Population recombination rate variation

We compared the broadscale (binned to 1 Mb) recombination rate variation of our Japanese genetic map and the HapMap (CEU + YRI + CHB/JPT; Utah residents (CEPH) with Northern and Western European ancestry; Yoruba in Ibadan, Nigeria; Han Chinese in Beijing, China / Japanese in Tokyo, Japan) populations (**Figure 1a, 1b**, and **Supplementary Fig. 2**). The broadscale recombination rate variation for the two maps was nearly identical. Both maps had higher rates in the sub-telomeric and peri-centromeric regions on some chromosomes and lower rates in the other middle regions. The cosine similarity score—a resemblance measure for the two profiles ranging from –1.0 (dissimilar) to 1.0 (identical)—was as high as 0.984 across the autosomes (**Supplementary Table 2**).

**Figure 1.**
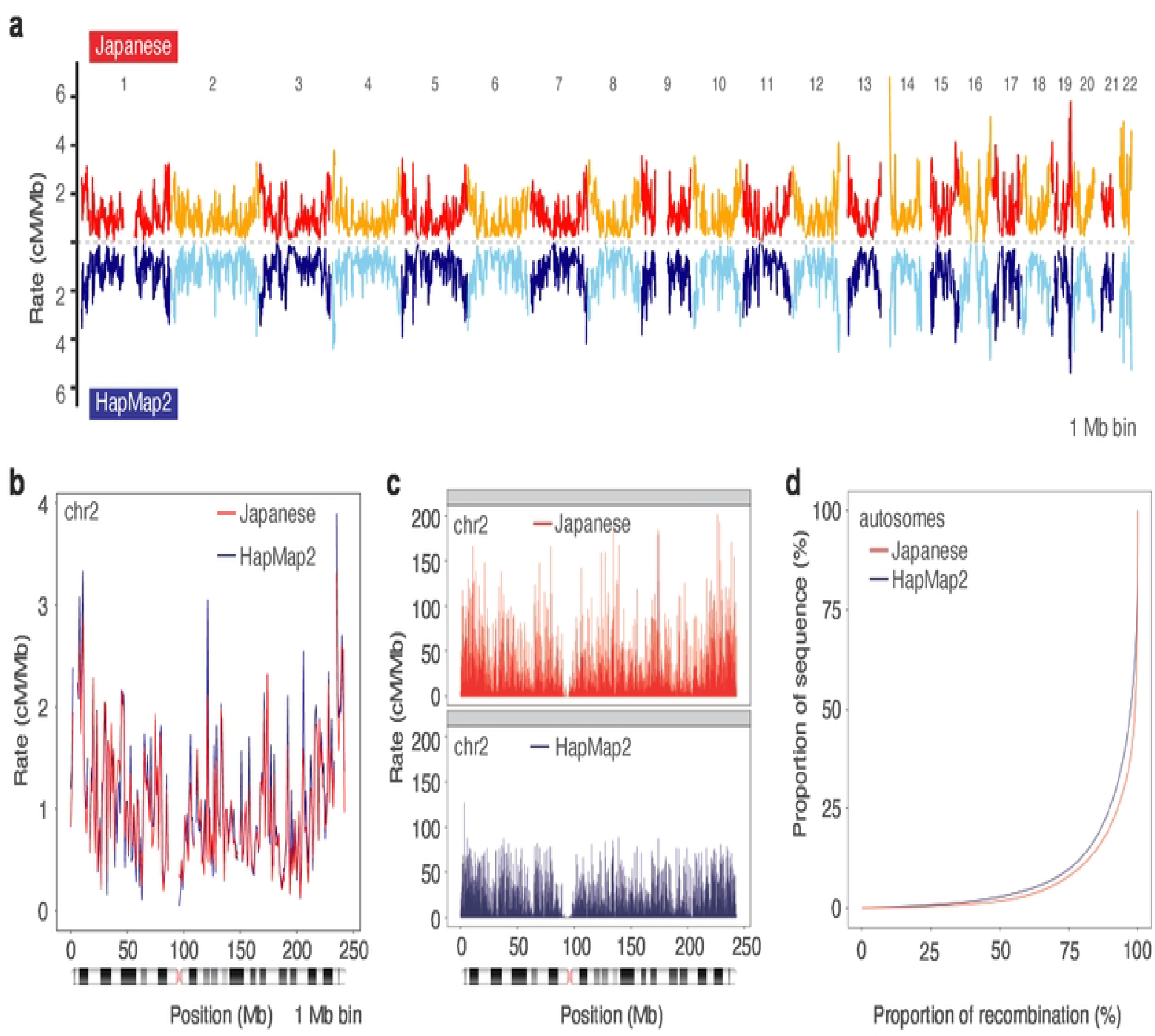
Linkage-disequilibrium-based genetic map for the Japanese population. **a** Broadscale concordance between the genetic map for the Japanese population and the HapMap2 (CEU + YRI + CHB/JPT) population. Rate, recombination rate. Numerals above the plot represent the chromosome (chr) number. **b** Concordance of the two genetic maps of chromosome 2, as an example. **c** Fine-scale recombination rate variation. Higher peaks were observed in our Japanese genetic map. **d** Recombination concentration analysis. The plot of proportion of sequence versus proportion of recombination showed more concentrated recombination in the Japanese population compared with the HapMap2 population.

The fine-scale recombination rate variation also resembled that of HapMap in terms of peak locations, but it differed in peak amplitude (**Figure 1c** and **Supplementary Fig. 3**). Our Japanese genetic map had 8970 sites with a per-generation-scaled recombination rate > 100 cM/Mb, whereas the HapMap genetic map had only one such site (**Figure 1c**).

These differences in the number and amplitude of the peaks may have been due to the finer resolution of our Japanese genetic map. Indeed, the number of SNP markers was 10,092,573 for the Japanese map and 3,303,922 for the HapMap genetic map across the genome. To evaluate the degree of recombination site concentration, we plotted the proportion of the sequence versus the proportion of recombination (**Figure 1d**). The Japanese recombination was more concentrated than that of the HapMap population.

For example, 90% of recombination occurred in 20.5% of the Japanese autosomes but in 25.5% of HapMap, and 80% of recombination occurred in 10.8% of the Japanese autosomes but in 13.0% of HapMap. This accumulation of recombination was plausibly due to the denser resolution of the Japanese genetic map.

### Recombination hotspots

From the recombination rate variation, we identified 9487 recombination hotspots with a maximum rate estimate ≥ 5 cM/Mb and size ≤ 5 kb (**Supplementary Table 3**). The average length of the hotspots was 3.9 kb after the filtering described above. The hotspot density ranged from 3.08 to 4.61 hotspots/Mb, and the average density was 3.53 hotspots/Mb across the autosomes. The number of hotspots from the HapMap genetic map was 34,012—much larger than the number from our map. This discrepancy was likely due to differences in the detection algorithms, filtering conditions, and definition of hotspots used, all of which make direct comparison difficult. More recent estimates using a software version close to our method have shown more similar values, namely 11,910 and 15,471 hotspots for European and African populations, respectively [12].

Recombination hotspots are associated with several sequence characteristics, such as the presence of a specific class of long terminal repeat transposon sequences called THE1A and THE1B, high GC%, and specific recombinogenic motifs, namely the 7-bp core motif, CCTCCCT [33], and the 13-bp extended motif, CCNCCNTNNCCNC [34]. To examine whether the 9487 hotspots had these characteristics, we prepared 10 control sets of 9487 randomly selected sequences of matched lengths from matched chromosomes. We confirmed the THE1A/B enrichment in the hotspots (**Table 1**). We also examined other classes of repetitive sequences and found results consistent with previous findings (**Table 1**). The GC% was higher in the hotspots than in the controls (42.5% vs. 41.0% ± 0.06%). Next, we examined whether the 7-bp and 13-bp motifs were enriched in the 9487 hotspots (**Figure 2**). We found that both motifs were significantly enriched (*P* = 8.33 × 10^−45^ to 2.69 × 10^−59^ and *P* = 7.4×10^−69^ to 2.1 × 10^−55^ for the 7-bp and 13-bp motifs, respectively; Fisher’s exact test with Bonferroni correction), and both motifs were localized at the center of the hotspots (*P* = 2.5 × 10^−65^ and *P* = 1.7 × 10^−19^ for the 7-bp and 13-bp motif, respectively; Fisher’s exact test with Bonferroni correction).

**Figure 2.**
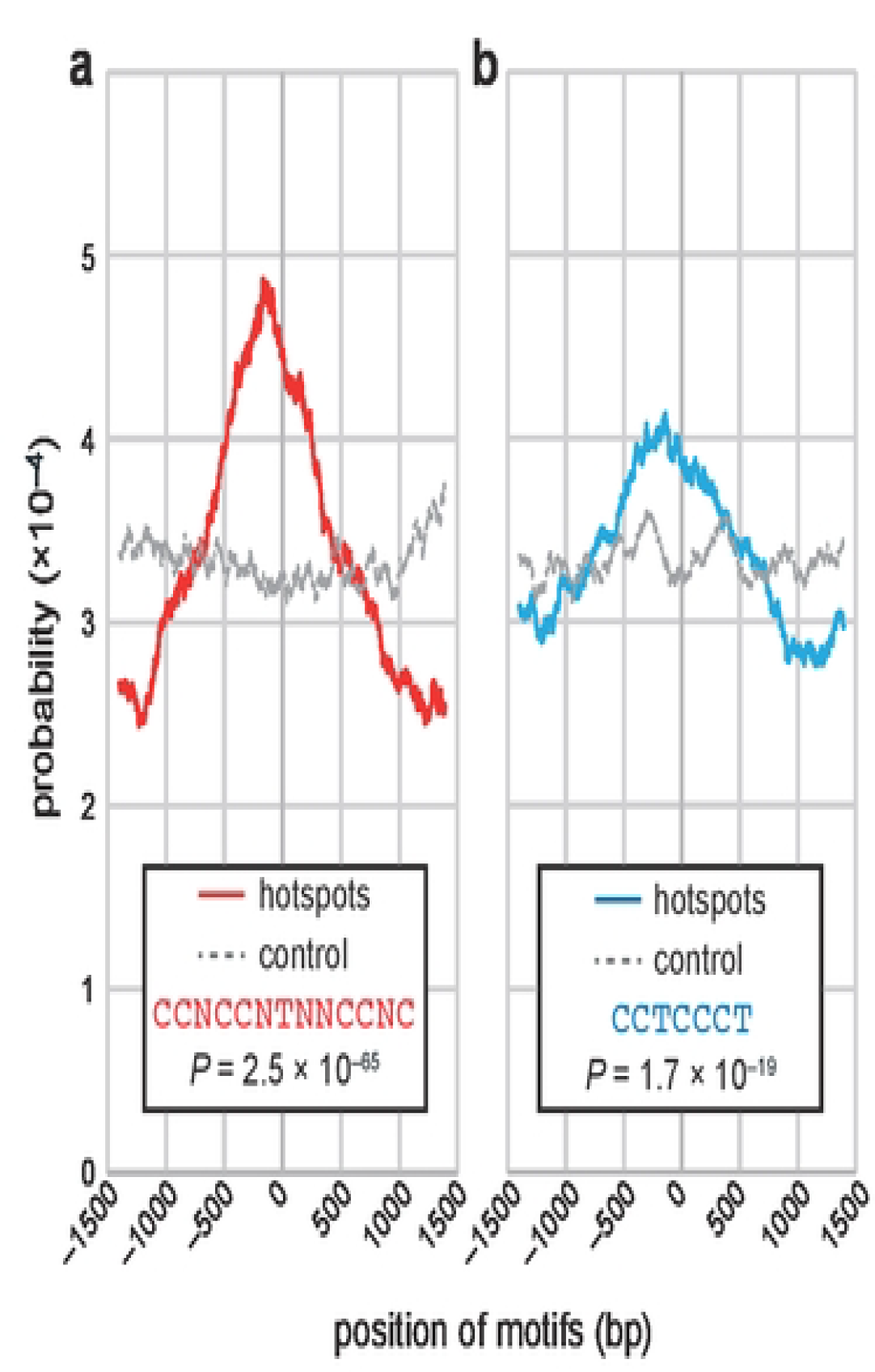
Enrichment of known motifs in hotspots. The 13-bp extended motif (**a**) and the 7-bp core motif (**b**) were enriched in the central regions of the identified hotspots. Solid lines in red or blue indicate enrichment in the hotspots, whereas dotted lines in gray indicate that in controls. Data were subjected to 500-bp-window moving average smoothing.

### Haplotype phasing and genotype imputation by using the genetic map

To examine the usefulness of the Japanese genetic map, we performed haplotype phasing using our genetic map and examined the accuracy. We used the WGS data of a cohort of 104 individuals of generation II from the TMM BirThree cohort, and performed phasing by using the Japanese genetic map or the 1KGP phase3 genetic map. The individuals used for the verification did not overlap with those used for the construction of the genetic mapping. The ground truth dataset was generated by performing phase-by-transmission with the parents’ WGS data. Then, we measured the switch error rate under 12 conditions. We found that the switch error rate was slightly lower with the Japanese genetic map in 10/12 conditions (**Table 2**).

We also performed a genotype imputation by using the Japanese genetic map and examined the accuracy. A cohort of 4768 Japanese subjects from the TMM BirThree cohort [17] was genotyped with SNP arrays. The 4768 samples were confirmed to be unrelated to each other and unrelated to the samples subjected to the genetic map construction. Their genotypes were then imputed with the Japanese haplotype reference panel (3.5KJPNv2) [18] by using either the Japanese genetic map or a conventional (1KGP) genetic map. We compared the INFO scores for each MAF (minor allele frequency) range in the 3.5KJPNv2 reference panel (**Table 3**). We found that the INFO scores in the imputed variants were higher when we used the Japanese genetic map (0.5471 ± 0.3739 vs. 0.5480 ± 0.3745 for conventional vs. Japanese map; n = 52,878,913 intersected variants; *P* = 1.46 × 10^−37^; Welch’s *t* test). Moreover, the INFO score improved for each MAF range (**Table 3**). Moreover, the number of sites with INFO scores above or equal to 0.8 increased from 19,934,605 to 20,042,021 when we changed from the conventional genetic map to the Japanese genetic map. Furthermore, among the 52,878,913 imputed variant sites, the INFO score increased for 47% of the sites, remained the same for 13%, and decreased for 40% when we used the Japanese genetic map. These results suggest that the use of population-specific genetic maps can improve the imputation accuracy for specific populations.

## DISCUSSION

We constructed a fine-scale genetic map for the Japanese population. The genetic map was constructed from 300 haploid genomes and 10 million SNP markers, the quality of which was carefully controlled, especially by the use of pedigree information. Pedigree information was also used for accurate haplotype phasing. The broadscale recombination rate variation was concordant with that of the HapMap genetic map, whereas the resolution was much improved. Moreover, across the autosomes, we identified 9487 hotspots that harbored sequence characteristics concordant with those in previous studies. We also demonstrated the use of the genetic map for genotype imputation.

Pedigree information, including genotype QC and haplotype phasing data, played a pivotal role in this study. Recombination rate variation can be inferred, in principle, from the genotypes of unrelated individuals [31]. However, because genotyping errors can mimic mutations and, in some cases, recombination events, genetic marker QC is critical for precisely inferring historical recombination events. Because the mutation rate for SNVs is substantially low (of the order of 10^−8^ per site per generation), most genotype calls inconsistent with Mendelian inheritance are presumably errors. We utilized the genotypes of parents and offspring, as determined by using the same experimental and bioinformatics pipelines for each, to keep the genotype quality high. Similarly, haplotype phasing is less prone to switch errors when direct relatives’ information is used than when a haplotype reference panel is used. Therefore, our strategy is suited to constructing high-quality genetic maps.

Population-specific genetic maps, along with other genetic resources, are set to become important population-specific genetic resources. To date, large-scale human genetic resources have been heavily biased toward European populations, with only a few analyses of other populations—especially Asian ones [35]. To address this deficiency, Asian population-specific genetic resources, such as haplotype reference panels [18, 36, 37] and reference genome sequences [38–41], are in active development. Although the improvement in imputation accuracy brought about by the use of our Japanese genetic map was minimal, genetic maps can be used for various genetic analyses, such as linkage mapping, haplotype phasing, genotype imputation, and genome-wide association studies. Therefore, the creation of population-specific genetic maps will help to establish more accurate statistical genetic analyses of underrepresented populations.

## DATA AVAILABILITY

The Japanese genetic map is available from the jMorp website (https://jmorp.megabank.tohoku.ac.jp/downloads/#genetic_map).

## ACKNOWLEDGMENTS

We thank S. Sugimoto for her help. This work was supported in part by the Tohoku Medical Megabank (TMM) Project of the Ministry of Education, Culture, Sports, Science and Technology and the Reconstruction Agency; and by the Japan Agency for Medical Research and Development (AMED; Grant Numbers JP20km0105001 and JP20km0105002) of Tohoku University. This work was also supported in part by JSPS KAKENHI Grant Numbers JP19H05200 to GT and JP19K06625 to JT. All computational resources were provided by the ToMMo supercomputer system (http://sc.megabank.tohoku.ac.jp/en), which is supported by the Facilitation of R&D Platform for AMED Genome Medicine Support, conducted by AMED (Grant Number JP20km0405001). We thank all the volunteers who participated in the TMM project.

## Notes

**Conflict of Interest:** The authors declare no conflict of interest.

### Competing Interest Statement

The authors have declared no competing interest.

## REFERENCES

1. Dib C, Fauré S, Fizames C, Samson D, Drouot N, et al. A comprehensive genetic map of the human genome based on 5,264 microsatellites. Nature. 1996;380(6570):152–154. doi: 10.1038/380152a0.

2. Broman KW, Murray JC, Sheffield VC, White RL, Weber JL. Comprehensive human genetic maps: individual and sex-specific variation in recombination. Am J Hum Genet. 1998;63(3):861–869. doi: 10.1086/302011.

3. Kong A, Gudbjartsson DF, Sainz J, Jonsdottir GM, Gudjonsson SA, Richardsson B, et al. A high-resolution recombination map of the human genome. Nat Genet. 2002;31:241–247.

4. Lichten M, Goldman ASH. Meiotic recombination hotspots. Annu Rev Genetics 1995;29:423–444.

5. Jeffreys AJ, Kauppi L, Neumann R. Intensely punctate meiotic recombination in the class II region of the major histocompatibility complex. Nat Genet. 2001;29:217–222.

6. Daly MJ, Rioux JD, Schaffner SF, Thomas J, Hudson TJ, Lander, ES. High-resolution haplotype structure in the human genome. Nat Genet. 2001;29:229–232.

7. Reich DE, Cargill M, Bolk S, Ireland J, Sabeti PC. et al. Linkage disequilibrium in the human genome. Nature 2001; 411, 199–204.

8. Stephens JC, Schneider JA, Tanguay DA, Choi J, Acharya T et al. Haplotype variation and linkage disequilibrium in 313 human genes. Science; 2001; 293, 489–493.

9. Gabriel SB, Schaffner SF, Nguyen H, Moore JM, Roy J et al. The structure of haplotype blocks in the human genome. Science. 2002;296(5576):2225–2229. doi: 10.1126/science.1069424.

10. McVean G, Awadalla P, Fearnhead PA. Coalescent-based method for detecting and estimating recombination from gene sequences. Genetics 2002;160:1231–1241.

11. McVean GAT, Myers SR, Hunt S, Deloukas P, Bentley DR, Donnelly P. The finescale structure of recombination rate variation in the human genome. Science 2004;304:581–584.

12. Auton A, Fledel-Alon A, Pfeifer S, Venn O, Ségurel L, Street T, et al. A fine-scale chimpanzee genetic map from population sequencing. Science 2012;336:193–198.

13. Slatkin M. Linkage disequilibrium—understanding the evolutionary past and mapping the medical future. Nat Rev Genet. 2008;9:477–485.

14. Service S, DeYoung J, Karayiorgou M, Roos JL, Pretorious H, Bedoya G, Ospina J, Ruiz-Linares A, Macedo A, Palha JA, Heutink P, Aulchenko Y et al. Magnitude and distribution of linkage disequilibrium in population isolates and implications for genome-wide association studies. Nat Genet. 2006;38(5):556–560. doi: 10.1038/ng1770.

15. The International HapMap Consortium. A second generation human haplotype map of over 3.1 million SNPs. Nature 2007;449:851–861.

16. The 1000 Genomes Project Consortium. A map of human genome variation from population scale sequencing. Nature 2010;467:1061–1073.

17. Kuriyama S, Metoki H, Kikuya M, Obara T, Ishikuro M, Yamanaka C, et al. Cohort profile: Tohoku Medical Megabank Project Birth and Three-Generation cohort study (TMM BirThree cohort study): rationale, progress and perspective. Int J Epidemiol. 2020;49:18–19m.

18. Tadaka S, Katsuoka F, Ueki M, Kojima K, Makino S, Saito S, et al. 3.5KJPNv2: an allele frequency panel of 3552 Japanese individuals including the X chromosome. Hum Genome Var. 2019;6:28.

19. O’Connell J, Gurdasani D, Delaneau O, Pirastu N, Ulivi S, Cocca M, et al. A general approach for haplotype phasing across the full spectrum of relatedness. PLoS Genet. 2014;10:e1004234.

20. The International SNP Map Working Group. A map of human genome sequence variation containing 1.42 million single nucleotide polymorphisms. Nature 2001;409:928–933.

21. Auton A, Myers S, McVean G. Identifying recombination hotspots using population genetic data. arXiv 1403.4264.

22. Quinlan AR, Hall IM. BEDTools: A flexible suite of utilities for comparing genomic features. Bioinformatics 2010;26:841–842.

23. Smit, AFA, Hubley, R & Green, P. RepeatMasker Open-4.0. 2013-2015< http://www.repeatmasker.org>.

24. McLeay R, Bailey TL. Motif Enrichment Analysis: A unified framework and method evaluation. BMC Bioinformatics 2010;11:165.

25. Bailey TL, Machanick P. Inferring direct DNA binding from ChIP-seq. Nucleic Acids Res. 2012;40:e128.

26. Delaneau, O., Zagury, JF. Marchini, J. Improved whole-chromosome phasing for disease and population genetic studies. Nat Methods 2013; 10, 5–6. 10.1038/nmeth.2307

27. Hofmeister, RJ., Ribeiro, DM., Rubinacci, S., Delaneau O. Accurate rare variant phasing of whole-genome and whole-exome sequencing data in the UK Biobank. BioRxiv 10.1101/2022.10.19.512867

28. Howie BN, Donnelly P, Marchini J. A flexible and accurate genotype imputation method for the next generation of genome-wide association studies. PLOS Genetics 2009; 5(6): e1000529. 10.1371/journal.pgen.1000529

29. Lee BT, Barber GP, Benet-Pagès A, Casper J, Clawson H, et al, The UCSC Genome Browser database: 2022 update, Nucleic Acids Res, 2022; 50 D1, (7) D1115–D1122, 10.1093/nar/gkab959

30. Sun, Y., Liu, F., Fan, C. et al. Characterizing sensitivity and coverage of clinical WGS as a diagnostic test for genetic disorders. BMC Med Genomics 2021; 14, 102. 10.1186/s12920-021-00948-5

31. Stumpf MPH, McVean GAT. Estimating recombination rates from population-genetic data. Nat Rev Genet. 2003;4:959–68.

32. Wang J, Raskin L, Samuels DC, Shyr Y, Guo Y. Genome measures used for quality control are dependent on gene function and ancestry. Bioinformatics 2015;31:318–323.

33. Myers S, Bottolo L, Freeman C. McVean G, Donnelly P. A fine-scale map of recombination rates and hotspots across the human genome. Science 2005;310:321–324.

34. Myers S, Freeman C, Auton A, Donnelly P, McVean G. A common sequence motif associated with recombination hot spots and genome instability in humans. Nat Genet. 2008;40:1124–1129.

35. Sirugo, G. Williams, S.M. Tishkoff S.A. The missing diversity in human genetic studies. Cell 2019;177:26–31.

36. Yoo SK, Kim CU, Kim HL, Kim S, Shin J-Y, Kim N, et al. NARD: whole-genome reference panel of 1779 Northeast Asians improves imputation accuracy of rare and low-frequency variants. Genome Med 2019;11:64.

37. GenomeAsia100K Consortium. The GenomeAsia 100K Project enables genetic discoveries across Asia. Nature 2019;576:106–111.

38. Cho Y, Kim H, Kim HM, Jho S, Jun JH, Lee YJ, et al. An ethnically relevant consensus Korean reference genome is a step towards personal reference genomes. Nat Commun 2016;7:13637.

39. Seo JS, Rhie A, Kim J et al., De novo assembly and phasing of a Korean human genome. Nature 2016;538:243–247.

40. Shi, L., Guo, Y., Dong, C. et al. Long-read sequencing and de novo assembly of a Chinese genome. Nat Commun 2016;7:12065. 10.1038/ncomms12065

41. Takayama J, Tadaka S, Yano K, Katsuoka F, Gocho C, Funayama T, et al. Construction and integration of three de novo Japanese human genome assemblies toward a population-specific reference. Nat Commun 2021;12:226.

